# Genetically engineered coral: A mixed-methods analysis of initial public opinion

**DOI:** 10.1101/2021.05.24.445411

**Authors:** Elizabeth V. Hobman, Aditi Mankad, Lucy Carter, Chantale Ruttley, Aditi Mankad, Lucy Carter, Chantale Ruttley

## Abstract

Rising sea surface water temperatures is contributing to coral degradation in the Great Barrier Reef. Synthetic biology technologies offer the potential to enhance coral resilience to higher water temperatures. To explore what the public think of genetically engineered coral, qualitative responses to an open-ended question in a survey of 1,148 of the Australian public were analysed. More respondents supported the technology (59%) than did not (11%). However, a considerable proportion indicated moderate or neutral support (29%). Participants commented about the (moral) right to interfere with nature and uncertainty regarding the consequences of implementing the technology. Participants also mentioned the need to take responsibility and act to save the reef, as well as the benefits likely to result from implementing the technology. Other themes included a desire for further testing and proof, more information, and tight regulation and controls when introducing the technology.

## 1. Introduction

The Great Barrier Reef (GBR) in Australia holds both significant and economic value (Great Barrier Reef Marine Park Authority, 2019). Home to more than 1,200 species of hard and soft corals (GBRMPA, 2019), the GBR is estimated to directly and indirectly contribute AU$11.6 billion to the Australian economy each year (Butler et al., 2013). Further, the GBR is integral to Australian identity, with 85% of Australians proud of its heritage status (Marshall, Curnock, Pert & Williams, 2019). It also holds significant heritage value to the traditional owners of the reef – the Aboriginal and Torres Strait Islander peoples (Australian Government, 2014).

### 1.1. Coral bleaching

Over the past five years, unprecedented coral loss has occurred as a result of rising seawater temperatures and increasing seawater acidity, crown-of-thorn starfish predation, poor water quality due to land-based sediment/nutrient/contaminant run-off, and localised cyclone damage (GBRMPA, 2019; Reef 2050 Water Quality Improvement Plan 2017-2022, 2018). Significant coral loss has been estimated across the GBR, especially in the northern region with an estimated 65% decline in coral cover since 2013 and an observed 10% coral cover in 2017, the lowest coral cover in 30 years of monitoring (Australian Government, 2014). This level of coral loss reflects a widespread threat to more than 3,000 reefs that make up the GBR and the marine ecosystems that these reefs support (Sheppard, Davy, Pilling, & Graham, 2018). While the Reef shows resilience in recovering after disturbances, the long-term trend for the GBR is one of decline due to the predicted increase in the frequency of chronic and acute disturbances, including coral bleaching events (Hughes et al., 2017, 2018, 2019). Although many factors play a role in coral bleaching, increasing sea surface temperatures associated with climate change present the most immediate threat to the reef (GBRMPA, 2019). Since 1994, annual sea surface temperatures have been above average (Bureau of Meteorology, 2018) and it is predicted that the chance of a marine heatwave will double that of today’s likelihood, if global mean temperatures reach the predicted 1.5 degrees (King, Karoly & Henley, 2017; Intergovernmental Panel on Climate Change, 2018; Henley & King, 2017; Leach, Millar, Haustein, Jenkins, Graham & Allen, 2018).

### 1.2. Current and emerging mitigating strategies

Effective reef management has mitigated some of the impacts of coral degradation on the GBR. These practices include initiatives like no-take (i.e., no removal of any sea flora or creatures) and no-entry zones (i.e., no human access), which have improved the reef’s ecosystems and resilience (McCook et al., 2010). However, given that bleaching impacts coral at a cellular level and its primary cause is increasing sea temperatures, solutions should also target this level of the organism and not just the surrounding physical environment (Anthony et al., 2017).

Synthetic biology is a new and emerging area of research that offers a potential suite of solutions to mitigate some of the negative impacts that coral reefs face due to environmental and biological factors. Rather than targeting extraneous factors contributing to coral reef degradation, synthetic biology can re-design DNA structures of the coral itself, making it more resilient to threats, such as increasing water temperatures (i.e., ‘heat resistant coral’). Specifically, the synthetic biology technology identifies natural gene variants in existing coral that enhance their ability to withstand higher temperatures and introduces these into corals to make them more heat resistant. Public acceptability of this type of new intervention, remains to be empirically tested.

### 1.3. Public perception of the reef

Prior research has shown that the public are concerned about coral reef degradation and feel the need to act urgently. For example, tourists visiting the GBR after a widespread coral bleaching event in 2017 reported more negatively-valenced emotional words when asked “*think about the first words that come to mind when you think of the Reef*”, compared to those visiting prior to the event (Curnock, Marshall, Thiault, Heron, Hoey, Williams, Taylor, Pert & Goldberg, 2017). By contrast, positively-valenced and neutral emotional words did not differ before and after the 2017 event. These results suggest that the public are becoming increasingly aware of the degradation to the GBR and are potentially experiencing what is known as ‘ecological grief’ (i.e., “the grief felt in relation to…ecological losses…due to…environmental change” Cunsolo & Ellis, 2018. While these negative emotions increased over time, it was also observed that ratings of value associated with the GBR (i.e., biodiversity value, scientific and educational value, lifestyle value, and international icon value), pride in the GBR, and GBR identity all increased from 2013 to 2017. There was also a concomitant increase in willingness to act and learn, despite decreases in personal responsibility, sense of agency, self-efficacy and optimism.

Although our study is exploratory and focuses on participants’ reactions to the genetic engineering of coral rather than the GBR itself, we anticipate that the public may provide responses that are similarly emotive, communicate the value/benefits of the reef, and suggest the need to remedy or reverse coral degradation.

### 1.4. Public perception of new technology

Although synthetic biology has the potential to positively influence scientific advancements across multiple domains, public perceptions of emergent scientific developments have the potential to negatively impact the development, commercialisation and implementation of synthetic biology applications. For example, the legalisation of embryos for stem cell research was severely impacted by public opinion and only introduced after lengthy public debate (Horst, 2008). Similarly, genetically modified organisms (GMOs) have faced widespread public criticism (and subsequently negatively impacted acceptance), without the public fully understanding the nuances of the technology (Nielsen, 2003). Based on public demand, most countries in the European Union now require mandatory labelling of genetically modified (GM) foods, and consumer advocates openly oppose the use of biotechnology in crop production (Moon & Balasubramanian, 2016).

Understanding how people formulate attitudes about emerging technologies provides important information about how technologies could be designed and implemented such that the public feels sufficiently confident in accepting their use. Given the increasing severity and occurrence of coral bleaching – coupled with the Australian public’s strong affinity with the GBR (Australian Government, 2014) – it is essential that we understand whether and why the public would accept synthetic biology solutions, prior to it being released or even developed in the first instance. By exploring these issues as early as possible, it will enable researchers to prepare for, and manage, likely public concerns in a responsible and effective manner. Although the Australian public’s reaction to synthetic biology in coral has yet to be investigated, research on synthetic biology technologies in other domains provides some insight into potential influences.

While limited, most of the research on public perceptions of synthetic biology has been conducted mainly in Europe and the U.S. (for a brief review, see Jin, Clark, Kuznesof, Lin & Frewer, 2019). This research, conducted across varying synthetic biology technologies, suggests that there is greater public enthusiasm for technologies that clearly address important societal, medical, or sustainability needs (Pauwels, 2013). For example, technologies focused on human health development and/or improvement are viewed favourably, whereas technologies that bring back extinct animals or technologies used for recreational purposes (e.g., glowing fish) are viewed least favourably and are considered an unacceptable use of this technology (Funk & Hefferon, 2018). It would appear then that the mechanism or intended purpose of the technology is an important consideration for people.

Furthermore, people seem to be predominantly attuned to the potential benefits and risks to animals, humans, and the ecosystem; moral, emotional or value-related issues (e.g., unnaturalness, creating life, playing God) (Dragojlovic & Einsiedel, 2012); and regulatory/control aspects (Akin et al., 2017; Betten, Broerse & Kupper, 2018; Hart Research Associates, 2013; Mandel, Braman & Kahan, 2008). Value predispositions (i.e., religiosity and deference towards scientific authority) and trust in scientists also have been found to be significantly correlated with support for synthetic biology (Akin et al., 2017; Dragojlovic & Einsiedel, 2012). Focussing further on perceived risks, some research also has reported public concern regarding the potential for secondary use, misuse and/or unintended consequences of synthetic biology (Gibson et al., 2010), including bioterrorism, loss of biodiversity, or the evolution of more resilient pests (Hart Research Associates, 2013; Newson, 2015; Rogers, 2011).

Psychological theories have also been developed to formally model some of these potential causal influences on individuals’ support or acceptance of novel scientific innovations and technologies more generally. For instance, in the area of novel, sustainable energy technologies (e.g., wind farms, carbon capture and storage, hydrogen vehicles, nuclear energy), a comprehensive framework has been proposed to explain technology acceptance (Huijts, Molin & Steg, 2012). This framework amalgamates several psychological theories and draws upon empirical evidence to a range of influences on technology acceptance including but not limited to perceived costs, risks and benefits; social norms; attitudes and perceived behavioural control (the Theory of Planned Behaviour: Azjen, 1991), outcome efficacy, problem perception and personal norms or moral obligations (the Norm Activation Model: Schwartz, 1968; 1977), affective influences, including both positive and negative affect (Loewenstein & Lerner, 2003), trust in the regulators or owners of the technology (Midden & Huijts, 2009; Siegrist & Cvetkovich, 2000), and fairness, including fairness of decision processes, and the distribution of outcomes (constructs borrowed from the organisational justice literature: Colquitt, 2001 with empirical evidence from wind farm studies: Gross, 2007; Wolsink, 2005). Across different technological applications, empirical work reveals support for the explanatory value of these social/psychological constructs.

Although this psychological framework can be applied to any technology that promises benefits along with potential risks and costs (Huijts et al., 2012), in the current study, we undertook an exploratory assessment of self-generated reasons for public support (or lack thereof) for a synthetic biology *solution* (i.e. heat resistant coral) to the *problem* of coral loss due to climate-related factors. Once these factors are identified, it may be then possible to theorise and experimentally test the causal influences on technology acceptance in this sphere. Thus, the focus of this study was to investigate the public’s awareness of and support for an emerging synthetic biology technology that involves modifying coral to enhance its resistance to increasing water temperatures. It aimed to address the following questions:

- What potentially influences peoples’ decisions about the acceptability of developing genetically engineered coral?
- What specific concerns do people hold regarding genetically engineered coral?

## 2. Method

### 2.1. Participants

One thousand one hundred and forty-eight (*N*=1,148) members of the public participated in this study. Imposed quotas ensured that the sample was representative of the Australian population on age (18-24years: *n*=146; 25-34 years: *n*=182; 35-44 years: *n*=168, 45-54 years: *n*=217; 55-64 years: *n*=193; 65 years or over: *n*=242), gender (535 males, 610 females, 3 other), and state of residence. A range of educational and employment levels were represented.

### 2.2. Procedure

Participants were recruited via an external third-party research agency with each participant receiving a token incentive for participation. To participate in the study, respondents were required to be an Australian resident and over the age of 18 years. The study was conducted during a 3-week period from November to December, 2018. The research received ethics approval from the CSIRO Social and Interdisciplinary Science Human Research Ethics Committee.

A standard introductory email was sent to potential participants, inviting them to take part in an online survey. Once participants clicked on the link to the survey, an information page was displayed explaining the general purpose of the study and inviting individuals to participate in the survey. Those that agreed to participate provided consent by ticking a checkbox and continuing with the survey. Initial demographic information (gender, age, postcode, state of residence) was collected to achieve quotas, thereby ensuring a representative sample of the Australian population on age, gender and location.

Towards the start of the survey, participants were provided with the following definition of synthetic biology:

- Synthetic biology is a new field of research bringing together genetics, chemistry and engineering. It allows scientists to design and build new biological organisms, so that they may perform new functions.
- Synthetic biology can use DNA^1^ to create new characteristics, or remove certain functions, in plants, animals and other organisms (e.g., bacteria, fungi, algae).

Participants then received information on the problem of coral loss in the GBR and a possible synthetic biology solution (i.e. genetic engineering of heat resistant coral). A power-point style presentation, or ‘storyboard’, was presented to participants to convey this information; this storyboard also provided textual and visual information about the novel technological solution. The storyboard was developed by the authors in collaboration with the scientists who are developing the technology, as well as CSIRO communication specialists. It should be noted that the information on the technology focussed on the benefits and did not explicitly detail potential negative consequences. The reasons for only presenting the benefits and not the negative consequences were two-fold – first, we wanted to see if and what negative outcomes people would raise themselves; and second, the technology is currently at such an infant stage of development that the potential negative outcomes are unknown at this stage.

Before viewing the storyboard, participants were asked whether they had heard of gene editing before. For those who had heard of it before, a knowledge of gene editing of coral was assessed by asking this subset of participants: “How much would you say you know about gene editing of coral?” (1=no knowledge, 2=a little knowledge, 3=some knowledge, 4=a lot of knowledge, 5=extensive knowledge).

Support for the coral technology was assessed by asking participants: “Overall, based on the information provided and your own general knowledge, to what extent would you support the development of this technology?” (1=would not support to 5=would strongly support).

After responding to this question, participants were asked the following questions: In deciding whether you’d support this technology, what influenced your decision? What is your main reason for supporting it, or not supporting it? An open text box was provided for participants to type their responses. They then completed a series of additional questions about the technology. Descriptive demographic information was requested at the end of the survey (e.g., education, and employment status). The survey took on average 15 minutes to complete.

### 2.3. Analytic approach

The qualitative analysis was conducted by the first author using the full data set (*n*=1148) and validity of coding was cross-checked by the third author (who coded 120 responses, for which kappa ranged from 0.88 to 1.00 across the final set of themes). A detailed coding scheme was developed iteratively for responses to the open-ended question: “In deciding whether you’d support this technology, what influenced your decision? What is your main reason for supporting it, nor not supporting it?”. To achieve parsimony and improve the meaningfulness of results, similar codes were combined prior to analysis. Only codes that were mentioned by at least 20 respondents were included in the final analysis. The codes were computed into several dummy-codes (0=did not mention this factor, 1=did mention this factor) and entered as predictors of support for the synthetic biology technology, in ordinary least squares regression. Note that the relative importance of the themes can be assessed by considering both: (a) the percentage of respondents who mentioned the theme and (b) the degree of change in support for the technology (as reflected in the unstandardized coefficients). In essence, important factors are those mentioned by a greater percentage of respondents and associated with a larger change in support ratings.

## 3. Results & Discussion

### 3.1. Knowledge of gene editing of coral

Most participants (n=811, 70.6%) had not heard of gene editing of coral before, indicating that awareness of this synthetic biology technology was relatively low. Proportionally more males were aware of the technology (n=182, 34.0%) than females (n=152, 24.9%) (χ^2^(1)=11.43, p=0.001). Age was significantly negatively correlated to awareness (*r*=−0.09, *p*=0.001) such that older people tended to be less aware. Education was not correlated to awareness (*r*=0.04, *p*=0.128).

Of the people who had heard of it before (n=337, 29.4%), self-reported knowledge was toward the lower end of the scale (Mean=2.06, SD=0.72). Less than 4% knew a lot or held extensive knowledge; the remainder held some or less knowledge. Thus, overall knowledge of genetically engineered coral was low across the sample.

These results mirror that of previous public attitude surveys in the U.S. showing that most people had heard nothing at all about synthetic biology (67% in 2008 to 43% in 2010: Hart Research Associates, 2013). While awareness appears to be increasing across the years (Pauwels, 2013), even if people have heard of the concept of synthetic biology, most do not feel generally well informed about it (Akin et al., 2017). For instance, in Australia in 2017, a little under half (48%) of the population had heard of synthetic biology but only 5% knew enough to explain it to a friend (Cormick & Mercer, 2017). Our study extends on this research by revealing that awareness and knowledge of a specific synthetic biology technology application – that is, genetically engineered coral – also is relatively low across the Australian population.

### 3.2. Support for gene editing of coral

Despite possessing low awareness and knowledge (prior to viewing information about the technology), participants generally supported the development of genetically engineered coral (Mean=3.69, SD=1.04, on a scale from 1=would not support to 5=would strongly support) (see Figure 2). Most people (~59%, *n*=682) supported the development of this technology by selecting either a ‘4’ or ‘5’ on the response scale. Approximately 11% (*n*=128) indicated less (selecting ‘2’) or no support (selecting ‘1’) for the development of heat tolerant coral. A good proportion (~30%) of people were moderately supportive (selecting ‘3’) – or, alternatively, undecided.

**Figure 1.**
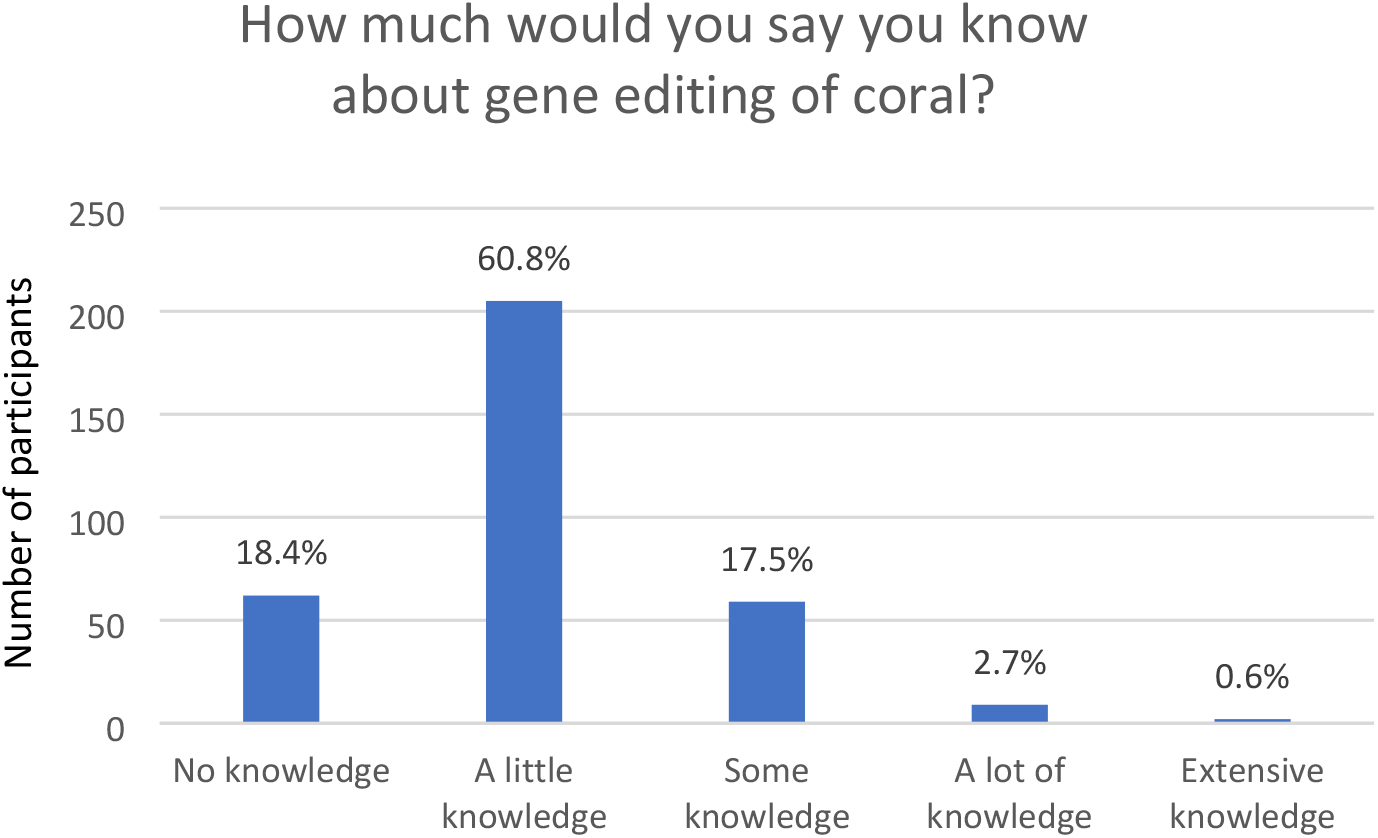
Knowledge of gene editing of coral (N=1,148)

**Figure 2.**
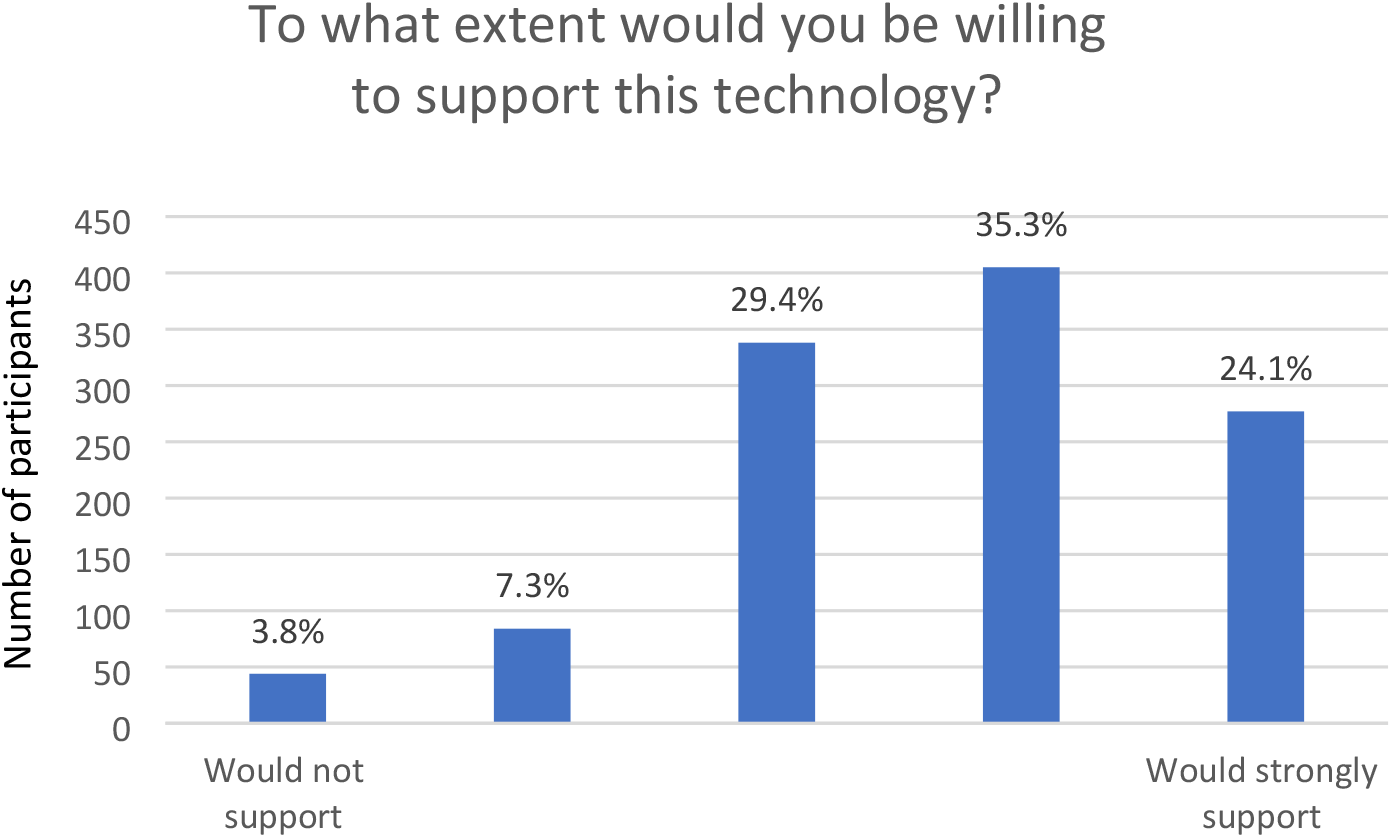
Support ratings for the development of the technology (N=1,148)

In terms of demographic correlates of support, age was significantly negatively related (*r*=−0.093, *p*=0.002) to support, as was sex (*r*=−0.075, p=0.012) – indicating that females were less supportive of the technology (Mean=3.61, SD=1.04) compared to men (Mean=3.77, SD=1.02). Educational level was not significantly related to support (*r*=0.03, *p*=0.269).

These results are somewhat consistent with previous quantitative and qualitative research in the U.S. where synthetic biology applications for environmental benefits are viewed quite favourably by the public (as are applications that address societal and medical needs; for an overview see Pauwels, 2013). Support for synthetic biology appears to strongly depend on its specific application (Ancillotti, Rerimassie, Seitz & Steurer, 2016; Funk & Hefferon, 2018). But even in the absence of an application, when a general definition of synthetic biology is provided (i.e., “Synthetic biology is the use of advanced science and engineering to make or redesign living organisms, such as bacteria. Synthetic biology involves making new genetic code, also known as DNA, that does not already exist in nature”: Akin et al., 2017, supplementary material, pg. 2), it has been observed that more people do not support its use (44%) than those that do (31%), with roughly a quarter ambivalent (i.e., neither supportive nor unsupportive) (Akin et al., 2017). Our results extend on these findings by indicating that many people are supportive of this specific application of synthetic biology in coral, however there is still a fair proportion who remain ambivalent or undecided. Further exploration of the self-professed reasons provided by participants may elucidate some of the concerns that people hold regarding the development of the technology.

### 3.3. Reasons for one’s support

A thematic analysis was conducted prior to a multiple regression analysis. This thematic analysis revealed several common themes. These themes were entered into a multiple regression analysis, whereby all themes together explained a significant amount of variance in support for the technology (*R*^2^= 0.49, F_(11,1137)_ = 100.42). Table 1 provides the unstandardized coefficients, beta coefficients and statistical significance. The table is ordered from most frequent to least frequently mentioned theme. Illustrative quotes for the themes are included both in the table and in the discussion below.

**Table 1.**
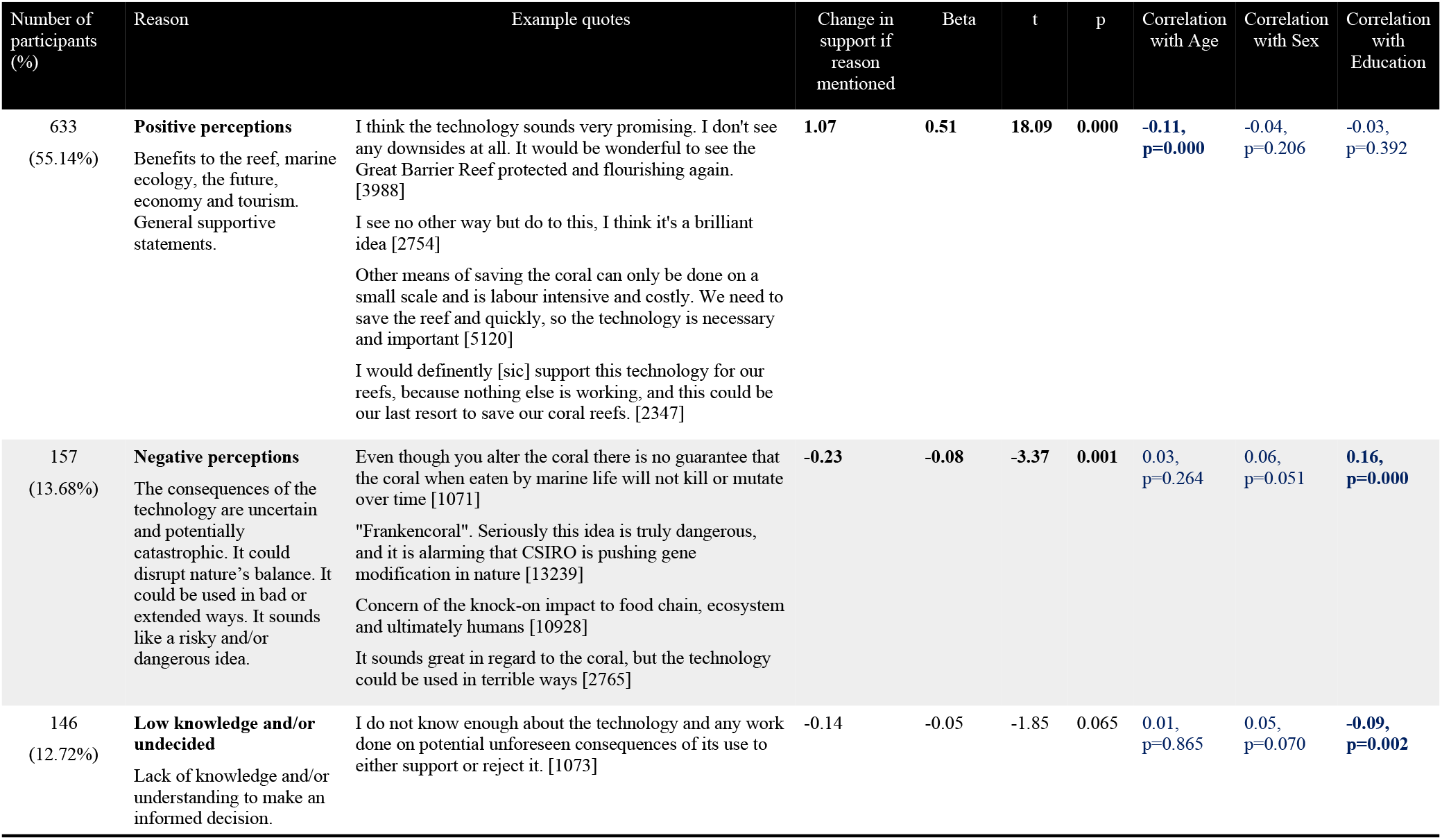
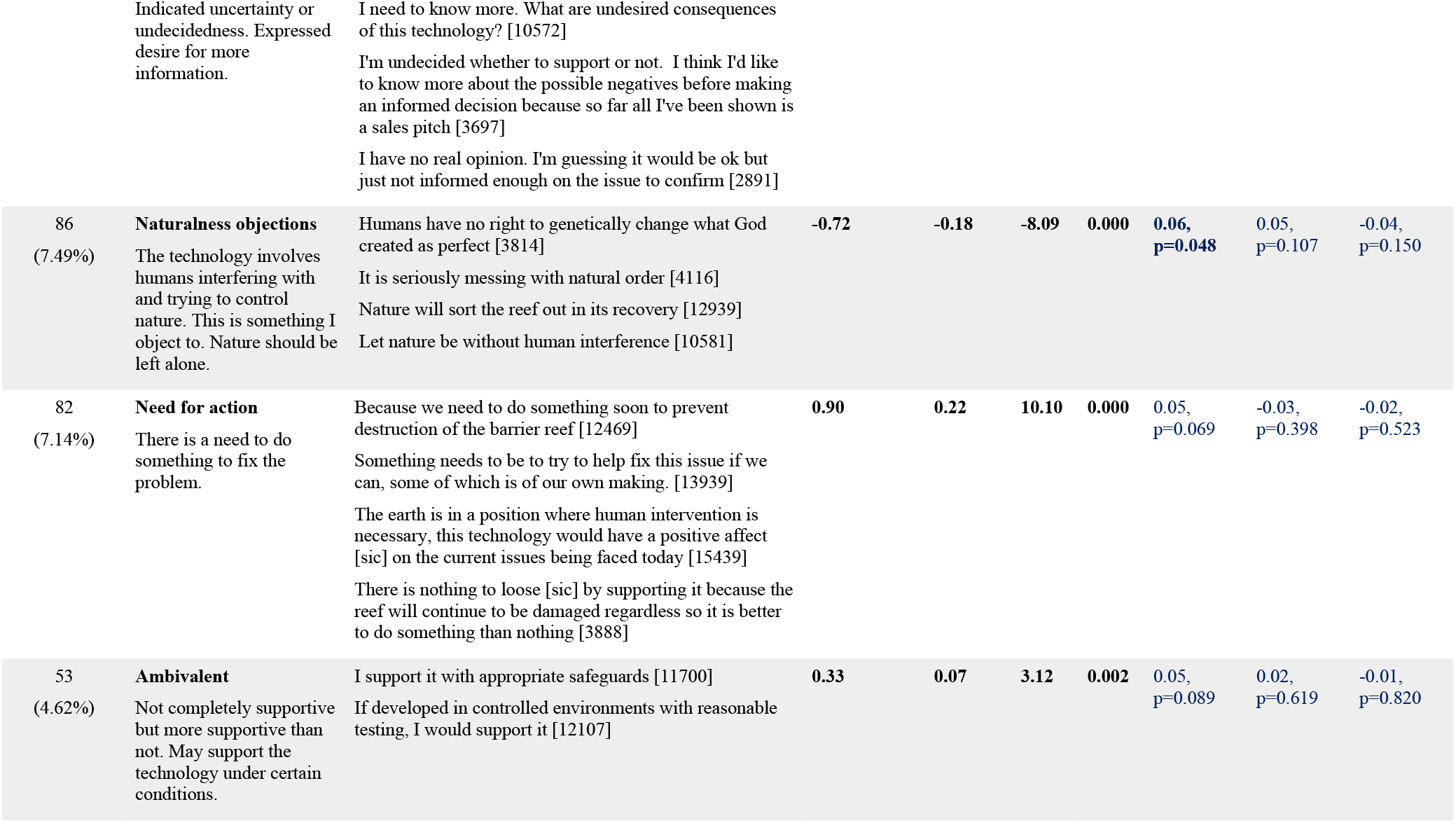
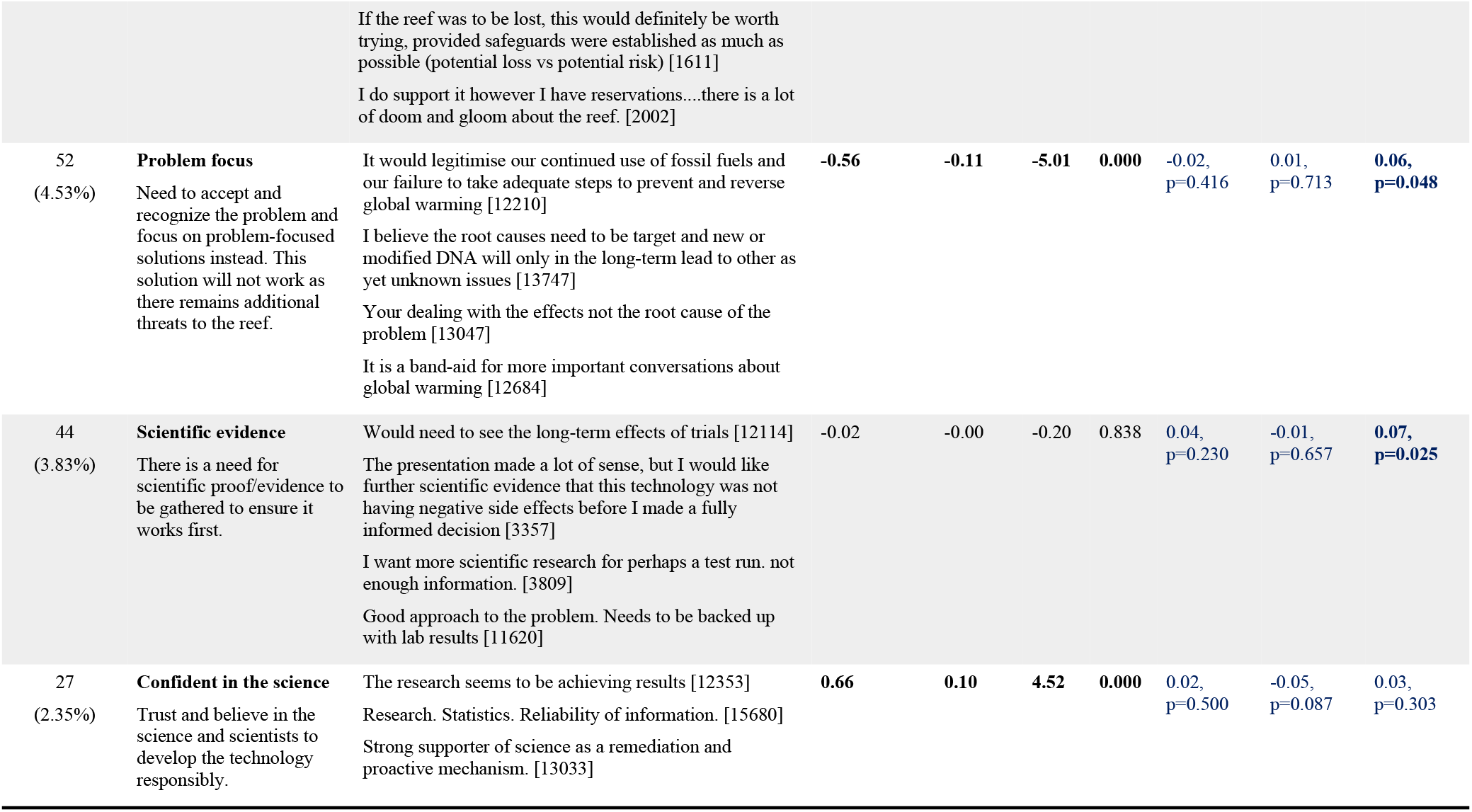
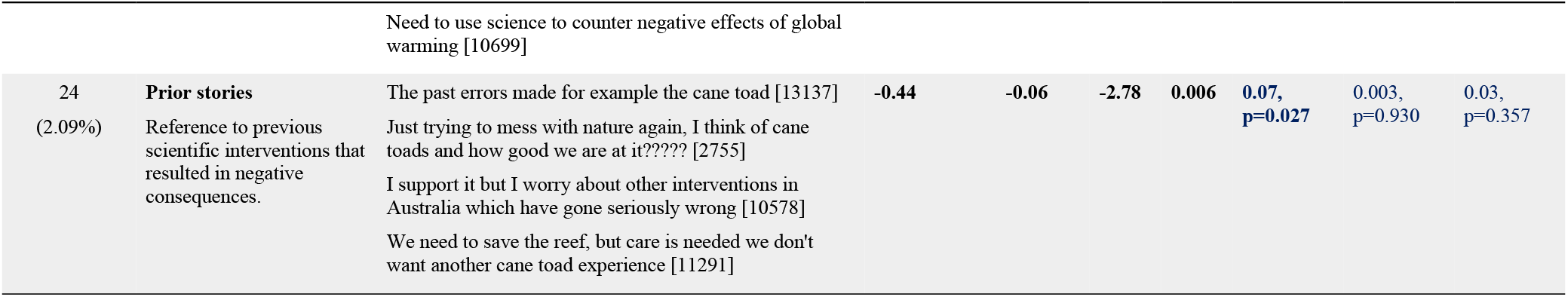
Common themes to explain one’s support for the development of the technology.

| Number of participants (%) | Reason | Example quotes | Change in support if reason mentioned | Beta | t | p | Correlation with Age | Correlation with Sex | Correlation with Education |
| --- | --- | --- | --- | --- | --- | --- | --- | --- | --- |
| 633<br>(55.14%) | **Positive perceptions**<br>Benefits to the reef, marine ecology, the future, economy and tourism. General supportive statements. | I think the technology sounds very promising. I don't see any downsides at all. It would be wonderful to see the Great Barrier Reef protected and flourishing again. [3988]<br>I see no other way but do to this, I think it's a brilliant idea [2754]<br>Other means of saving the coral can only be done on a small scale and is labour intensive and costly. We need to save the reef and quickly, so the technology is necessary and important [5120]<br>I would definitely [sic] support this technology for our reefs, because nothing else is working, and this could be our last resort to save our coral reefs. [2347] | **1.07** | **0.51** | **18.09** | **0.000** | **-0.11, p=0.000** | -0.04, p=0.206 | -0.03, p=0.392 |
| 157<br>(13.68%) | **Negative perceptions**<br>The consequences of the technology are uncertain and potentially catastrophic. It could disrupt nature's balance. It could be used in bad or extended ways. It sounds like a risky and/or dangerous idea. | Even though you alter the coral there is no guarantee that the coral when eaten by marine life will not kill or mutate over time [1071]<br>"Frankencoral". Seriously this idea is truly dangerous, and it is alarming that CSIRO is pushing gene modification in nature [13239]<br>Concern of the knock-on impact to food chain, ecosystem and ultimately humans [10928]<br>It sounds great in regard to the coral, but the technology could be used in terrible ways [2765] | **-0.23** | **-0.08** | **-3.37** | **0.001** | 0.03, p=0.264 | 0.06, p=0.051 | **0.16, p=0.000** |
| 146<br>(12.72%) | **Low knowledge and/or undecided**<br>Lack of knowledge and/or understanding to make an informed decision. | I do not know enough about the technology and any work done on potential unforeseen consequences of its use to either support or reject it. [1073] | -0.14 | -0.05 | -1.85 | 0.065 | 0.01, p=0.865 | 0.05, p=0.070 | **-0.09, p=0.002** |
|  | Indicated uncertainty or undecidedness. Expressed desire for more information. | I need to know more. What are undesired consequences of this technology? [10572]<br>I'm undecided whether to support or not. I think I'd like to know more about the possible negatives before making an informed decision because so far all I've been shown is a sales pitch [3697]<br>I have no real opinion. I'm guessing it would be ok but just not informed enough on the issue to confirm [2891] |  |  |  |  |  |  |  |
| 86<br>(7.49%) | **Naturalness objections**<br>The technology involves humans interfering with and trying to control nature. This is something I object to. Nature should be left alone. | Humans have no right to genetically change what God created as perfect [3814]<br>It is seriously messing with natural order [4116]<br>Nature will sort the reef out in its recovery [12939]<br>Let nature be without human interference [10581] | -0.72 | -0.18 | -8.09 | 0.000 | 0.06,<br>p=0.048 | 0.05,<br>p=0.107 | -0.04,<br>p=0.150 |
| 82<br>(7.14%) | **Need for action**<br>There is a need to do something to fix the problem. | Because we need to do something soon to prevent destruction of the barrier reef [12469]<br>Something needs to be to try to help fix this issue if we can, some of which is of our own making. [13939]<br>The earth is in a position where human intervention is necessary, this technology would have a positive affect [sic] on the current issues being faced today [15439]<br>There is nothing to loose [sic] by supporting it because the reef will continue to be damaged regardless so it is better to do something than nothing [3888] | 0.90 | 0.22 | 10.10 | 0.000 | 0.05,<br>p=0.069 | -0.03,<br>p=0.398 | -0.02,<br>p=0.523 |
| 53<br>(4.62%) | **Ambivalent**<br>Not completely supportive but more supportive than not. May support the technology under certain conditions. | I support it with appropriate safeguards [11700]<br>If developed in controlled environments with reasonable testing, I would support it [12107] | 0.33 | 0.07 | 3.12 | 0.002 | 0.05,<br>p=0.089 | 0.02,<br>p=0.619 | -0.01,<br>p=0.820 |
|  |  | If the reef was to be lost, this would definitely be worth trying, provided safeguards were established as much as possible (potential loss vs potential risk) [1611]<br>I do support it however I have reservations....there is a lot of doom and gloom about the reef. [2002] |  |  |  |  |  |  |  |
| 52<br>(4.53%) | **Problem focus**<br>Need to accept and recognize the problem and focus on problem-focused solutions instead. This solution will not work as there remains additional threats to the reef. | It would legitimise our continued use of fossil fuels and our failure to take adequate steps to prevent and reverse global warming [12210]<br>I believe the root causes need to be target and new or modified DNA will only in the long-term lead to other as yet unknown issues [13747]<br>Your dealing with the effects not the root cause of the problem [13047]<br>It is a band-aid for more important conversations about global warming [12684] | **-0.56** | **-0.11** | **-5.01** | **0.000** | -0.02,<br>p=0.416 | 0.01,<br>p=0.713 | **0.06,**<br>**p=0.048** |
| 44<br>(3.83%) | **Scientific evidence**<br>There is a need for scientific proof/evidence to be gathered to ensure it works first. | Would need to see the long-term effects of trials [12114]<br>The presentation made a lot of sense, but I would like further scientific evidence that this technology was not having negative side effects before I made a fully informed decision [3357]<br>I want more scientific research for perhaps a test run. not enough information. [3809]<br>Good approach to the problem. Needs to be backed up with lab results [11620] | -0.02 | -0.00 | -0.20 | 0.838 | 0.04,<br>p=0.230 | -0.01,<br>p=0.657 | **0.07,**<br>**p=0.025** |
| 27<br>(2.35%) | **Confident in the science**<br>Trust and believe in the science and scientists to develop the technology responsibly. | The research seems to be achieving results [12353]<br>Research. Statistics. Reliability of information. [15680]<br>Strong supporter of science as a remediation and proactive mechanism. [13033] | **0.66** | **0.10** | **4.52** | **0.000** | 0.02,<br>p=0.500 | -0.05,<br>p=0.087 | 0.03,<br>p=0.303 |
|  |  | Need to use science to counter negative effects of global warming [10699] |  |  |  |  |  |  |  |
| 24<br>(2.09%) | **Prior stories**<br>Reference to previous scientific interventions that resulted in negative consequences. | The past errors made for example the cane toad [13137]<br>Just trying to mess with nature again, I think of cane toads and how good we are at it???? [2755]<br>I support it but I worry about other interventions in Australia which have gone seriously wrong [10578]<br>We need to save the reef, but care is needed we don't want another cane toad experience [11291] | **-0.44** | **-0.06** | **-2.78** | **0.006** | **0.07, p=0.027** | 0.003, p=0.930 | 0.03, p=0.357 |

#### 3.3.6. Positive perceptions

Many (*n*=633, ~55%) participants made comments that could be classified under the broad theme of positive perceptions. This theme primarily centred on the benefits that the technology would bring to the reef in terms of saving the reef, the broader marine system and associated industries (e.g., tourism). Additionally, some participants simply provided a generalised positive statement about the technology, sometimes comparing it favourably to other solutions.

> *To help rebuild our reefs* [3350]

> *The importance of saving the Great Barrier Reef and its associated inhabitants* [6057]

> *Yes I would to help protect and restore the barrier reef because it is 1 of Australia’s wonders and to help the marine life* [3844]

Those who made a comment under this theme were significantly more likely to support the development of the technology (t=18.09, p=0.000). Prior qualitative research has revealed that people discuss synthetic biology in a positive light mainly by expressing hope that the technology could successfully address significant societal and environmental challenges (Bhattachary et al., 2010; van Mil et al., 2017). Quantitative estimates are that around 1 in 5 people (22%) agree that synthetic biology would yield high benefits (Akin et al., 2017). Thus, prior research suggests that people do recognise the potential benefits of synthetic biology, however, any optimism expressed is usually caveated with much caution and conditions, such as strong governance, transparency and information (Ancillotti et al., 2016; Bhattachary, Calitz & Hunter, 2010). In our study, we seemed to observe more widespread positive opinion, though this could be due to our focus on one specific technology, that is, genetically engineering coral – and the fact that responses were limited to a single comment box, reducing the potential for participants to elaborate on their initial points of view. Additionally, our storyboards intentionally only presented the potential positive outcomes of using the technology rather than providing a balanced set of information on the benefits and risks. If balanced information were to be provided, it is likely that a more nuanced comment/opinion would have resulted (Pauwels, 2013). Once the potential risks or negative consequences are known – as the technology develops – these factors could well be included in further public surveys and discussions.

#### 3.3.1 Negative perceptions

On the converse, around 14% (*n*=157) of the total sample mentioned reasons that reflected negative perceptions about the technology. Most comments referred to the risk of potential negative consequences. Examples include:

> *Even though you alter the coral there is no guarantee that the coral when eaten by marine life will not kill or mutate over time* [Participant 1071]

> *We do not know enough about the long-term possibilities* [Participant 3179]

> *Could create a bad coral strand, could become toxic destroying more than it is now* [Participant 10831]

These negative perceptions, while not necessarily widespread, were significantly associated with less support (t=−3.39, p=0.001). Participants either referred to how the consequences of the technology were uncertain and potentially negative (especially for the environment and broader ecosystem); the potential for the technology to be used in other, negative ways beyond its original application; or more generally, they simply stated that the technology sounded ‘risky’ or ‘dangerous’.

These results are consistent with the Theory of Planned Behaviour (Azjen, 1991) whereby people are thought to form attitudes based on an appraisal of the perceived costs/risk and benefits. In the context of synthetic biology, perceived risks and benefits is a salient and topical theme. For example, survey research in the U.S. has revealed that about a quarter think the risks will be high, while a similar percentage also believe the benefits will be high – and when a single ‘risks – benefits’ measure is created, it correlates negatively and significantly with support (Akin et al., 2017). Other research has shown that concerns about risks outweighing the benefits significantly heightens (i.e., doubling the percentage sharing this concern) when people are provided balanced information (including its potential benefits and risks) as opposed to when no information is provided (Pauwels, 2013). In focus groups where balanced information is provided, discussions appear to be more nuanced whereby participants express more ambivalence towards the technology (Pauwels, 2013). Along with perceived risks, the themes of uncontrollability, intended harm (via misuse/dual use) and unintended harm (i.e., accidents, unforeseen mutations/evolution) have been observed in European focus group/workshop discussions (Ancillotti et al., 2016; Bhattachary et al., 2010).

#### 3.3.2. Low knowledge and/or undecided

Reflecting the relatively low levels of awareness and knowledge of genetically engineering coral as measured quantitatively, many participants (*n*=146, 12.72%) explicitly stated that their current base of knowledge regarding the technology was insufficient for deciding whether to support the technology. They explained that their knowledge is lacking, they did not understand or know the technology very well, and required more information, especially about broader consequences of the technology and associated risks, in order to make an informed decision. Some also explicitly stated that they did not know, were unsure or undecided about the technology. All these reasons were combined under the broad theme of low knowledge and/or undecided.

> *I need to know more. What are undesired consequences of this technology?* [2612]

> *Need more information on long term impact to other marine wildlife* [2283]

> *Still find it hard to understand what it is all about* [2566]

While approaching significance, the results showed that those who mentioned that they did not have enough knowledge were no more or less likely to support the development of the technology (t=−1.85, p=0.065).

As explained earlier, awareness and/or knowledge of synthetic biology – let alone specific applications – is quite low among the general population, though awareness appears to be on the rise. In the context of this low knowledge and the fact that the specific gene editing technology is in its infancy, it is not surprising that people are seeking more information in our study. Our results are consistent with previous research revealing that people are interested in, and seeking more information about, synthetic biology; requesting this communication effort to be completely transparent, accessible and available to the public (Ancillotti et al., 2016; Pauwels, 2013; van Mil, Hopkins & Kinsella, 2017). In the absence of such knowledge, research suggests that people may more heavily rely on a variety of other cues to determine whether they will support the technology – factors such as deference to scientific authority and trust in scientists (Akin et al., 2017).

#### 3.3.3. Naturalness objections

A small number (*n*=86, ~7%) of people made comments that fundamentally objected to human interference in nature or more broadly how nature should be left alone presented. The language used in these comments reflected a firm stance. Examples include:

> *The reef should be allowed to go through its natural cycle* [1071]

> *Human interference in nature, although ostensibly for the good, has too often resulted in the creation of a new set of problems* [3205]

> *We should just leave nature as it is* [5092]

Of all the themes negatively related to support, naturalness objections were the most significant correlate (t=-8.09, p=0.000), highlighting its importance in potentially influencing one’s level of support.

Certainly, prior research has revealed that some people may not support synthetic biology due to an underlying yet highly accessible implicit belief that genetic manipulation is unnatural (and therefore morally wrong); or alternatively, they may hold a more explicit evaluation that synthetic biology violates God’s domain as the creator of life (Dragojlovic & Einsiedel, 2012). For example, it has been observed that those who are more religious (or who believe in God) tend to be more concerned about the risks of synthetic biology (Kahan, Braman & Mandel, 2009; Dragojlovic & Einsiedel, 2012; Hart Research, 2013) and are less supportive of using the technology (Akin et al., 2017; Dragojlovic & Einsiedel, 2012). Even among people who may be more moderate in their view of synthetic biology also raise concerns about man intervening in nature (Ancillotti et al., 2016; Bhattachary, 2010), and the morality of constructing life (Ancillotti et al., 2016). Yet other research has suggested these types of objections are used to convey unease with more tractable concepts such as the (potentially destructive) power of humans to alter entire ecosystems (National Academies of Sciences, Engineering, and Medicine, 2016), slippery slope arguments (Link 2013) and distrust in scientific advancements (Raimi, Wolske, Sol Hart & Campbell-Arvai, 2019). Together, these results suggest that for some individuals, value predispositions may play a significant role in determining how they view synthetic biology, potentially overshadowing an appraisal of risks and benefits.

#### 3.3.7. Need for action

A smaller proportion (*n*=82, ~7%) explained that it was important to act sooner rather than later; to do something to save the reef given the threat of its destruction, and the fact that humans are accountable/responsible for remedying the situation.

> *We need to keep on improving and finding ways to get it fix* [7000]

> *All avenues should be pursued to assist* [6149]

> *Humans aren’t doing enough to slow climate change, so whilst I would rather nature run it’s course, I can see the benefits of this technology and am afraid without it the coral reef will be lost* [3031]

Participants who mentioned this reasoning were significantly more likely to support the technology (t=10.10, p=0.000). Our findings resonate with Curnock and colleagues’ (2019) assessment that people (in this case, tourists) possess a strong and increasing desire to help protect the GBR despite feeling only moderately able to take individual action themselves. It is therefore not surprising that in the context of genetically engineering coral that some participants communicated the desire for ‘us’ humans, in the collective sense, to take action.

#### 3.3.8. Ambivalent

There was also a small proportion (~5%) who specifically explained that they were ambivalent in their support or that their **s**upport was conditional, such as wanting the technology to be developed in a safe and controlled manner, or to be sure that there would be no negative consequences.

> *If developed in controlled environments with reasonable testing, I would support it* [2015]

> *I do support it however I have reservations…there is a lot of doom and gloom about the reef* [2021]

> *If it helps to keep everything going and not die off then it’s a very good thing to be doing and also if it is all monitored with the experts, why not* [3037]

Participants who mentioned this reasoning were significantly more likely to support the technology (t=3.10, p=0.002). As shown by the representative comments, people tended to say that they did support the technology so long as there were some controls or safeguards in place. Ultimately, they were expressing some caution and reservations despite thinking positively towards the technology. As discussed earlier, this type of ‘cautious optimism’ is a commonly observed in previous qualitative research with the public (Ancillotti et al., 2016; Pauwels, 2013; van Mil et al., 2017).

#### 3.3.4. Problem focus

A small proportion (~5%) mentioned that the technology could or would subvert a more sensible problem focus (i.e., attending to the root causes of rising sea surface temperatures), and that the technology would be insufficient to save the Reef given the range of other threats facing the GBR (e.g., the crown-of-thorns starfish and cyclones). Those who believed there needed to be a focus on managing the root cause of the problem, and that the technological solution could be futile, tended to be less supportive of the technology overall (t=−5.01, p=0.000). Example comments included:

> *It is the role of humans to change their environment damaging behaviour and not changing natural organisms*… [3814]

> *Overpopulation, poor farming techniques, tackling climate change need to be addressed* [2728]

> *It appears to be putting resources into developing a highly sophisticated solution to a symptom of global warming, rather than focusing those resources on addressing the causes of global warming* [12118]

Consistent with our results, prior research has found that people are motivated to explore and consider alternative or additional ways of addressing societal and/or environmental problems, highlighting the complex systems in which problems occur, as well as the root causes to these problems (Ancillotti et al., 2016; Bhattachary et al., 2010; van Mil et al., 2017). Rather than viewing gene-based technologies as a panacea to complex problems caused by multiple and interrelated factors, these results suggest the public expect to engage with science differently. People are conscious of the need to consider the problem context more fully, to explore the utility of alternative solutions (including non-technological solutions such as behaviour change), and to consider using genetic solutions in concert with other tools/solutions, ultimately forming a ‘suite of solutions’ (van Mil et al., 2017).

#### 3.3.5. Scientific evidence

Separate to the *lack of knowledge* theme were comments that explicitly asked for scientific evidence or trials to be conducted to evaluate the technology. People wanted to see long-term trials and scientific evidence demonstrating that there were no negative side effects from introducing the technology.

> *I want more scientific research for perhaps a test run. Not enough information.* [2008]

> *Would need to see evidence of successful trials and consider any unexpected longer term consequences before deciding to fully support this concept* [3023]

> *Would need to see the long term effects of trials* [3039]

A small proportion (~4%) mentioned the need for scientific testing/evidence, however, it was not significantly related to whether they supported the technology (t=−0.20, p=0.838). Approximately 57% of people who mentioned this theme were neutral in their support of heat resistant coral, with ~14% and ~29% expressing less and more support, respectively. It is possible, then, that this small group of people were feeling somewhat cautious towards the technology, and therefore wanted to see more reliable evidence from scientific trials.

As discussed above, previous research exploring public perceptions of synthetic biology has revealed that many people are interested in learning more, especially in the context of having low base levels of knowledge about the technologies (Ancillotti et al., 2016; van Mil et al., 2017). Our findings provide nuances surrounding this desire for information by revealing that some people may want to observe the outcomes of scientific trials that demonstrate the efficacy of the technology (or otherwise). Again, this is consistent with the Theory of Planned Behaviour, which proposes that people form attitudes in part through an appraisal of the risks/costs and benefits.

#### 3.3.8. Confident in the science

Finally, there was a smaller proportion (~2%) again who explained that they had confidence in and trusted the science and evidence that the technology would indeed be an effective solution.

> *It seems it’s strongly based on scientific fact without damaging the ecological environment* [2077]

> *I’m confident it’s been tested vigorously in this day and age* [4350]

> *The fact that testing will be controlled in a lab situation before anything is done in the reef* [2663]

Participants who mentioned this reasoning were significantly more likely to support the technology (t=4.52, p=0.000). Prior research has revealed that scientists, academics and researchers are one of the most trusted actors to work on genetic technology (van Mil et al., 2017), and trust in scientists has been shown to be significantly and positively associated with support for synthetic biology (Akin et al., 2017). Reasons given for trusting scientists include their impartiality, independence, capabilities, competence and experience (van Mil et al., 2017). Our results slightly differ from this prior research in that our participants spoke not only about trust in the agency or organisation that would oversee its development (e.g., CSIRO), but they also spoke about trust and confidence in the actual conduct of the science and research itself. It is possible that people would have commented about trust (or otherwise) in the scientists developing the technology, if scientists had been a focus of presentation in the storyboard.

#### 3.3.6. Prior stories

The final theme that was significantly negatively related to support was prior stories of failed scientific field interventions, such as the introduction of the cane toad into Australia. While only a small percentage (*n*=24, ~2%) of participants mentioned previous stories, it was still significantly related to less support for the technology, signifying how salient historical failures of introduced bio-based technologies are in the minds of some (t=−2.78, p=0.006). Examples include:

> *Just trying to mess with nature again, I think of cane toads and how good we are at it????* [5042]

> *The past errors made for example the cane toad* [60004]

> *Don’t know if this could be detrimental in the long run – as has happened with Monsanto interfering with the growing of cereal crops* [4702]

As observed in previous qualitative research (Ancillotti et al., 2016), our participants drew links between the current synthetic biology technology and past experiences or stories that readily came to mind. It is possible that both the availability heuristic (Tversky & Kahneman, 1973) and the negativity bias (Kanouse & Hanson, 1971) are in operation here. The availability heuristic is a mental short-cut that involves making judgements or decisions based on how easily an example, story or similar experience comes to mind – that is, when an example spring to mind quickly, people may perceive that example (and the information it conveys) as important and more heavily rely on this example to assist them in forming a decision. While this process certainly improves the speed of decision-making, in some cases, it can lead to biased decisions. Further to this, the negativity bias explains that more negative things (e.g., unpleasant thoughts, emotions, events) have a greater effect on one’s psychological state than do neutral or positive things (Baumeister, Finkenauer & Vohs, 2001). Applied to memory, research reveals that people tend to recall negative events/information more so than positive events/information (Finkenauer & Rime, 1998; Robinson-Riegler & Winton, 1996; Thomas & Diener, 1990). Thus, in the case of remembering negative stories such as the introduction of the cane toad, people would then be more likely to consider that the current technology could similarly fail too.

## Conclusions & Implications for Technology Development

This study has identified a range of factors that may explain support (or otherwise) for the development of genetically engineered coral. Many people felt that the technology would yield important benefits for the reef and that there were no other viable alternatives. Some also raised the point that humans are responsible for taking action to remedy the problem that humans as a collective group have contributed to. While a few people expressed confidence in the science underpinning the technology, other people exhibited more ambivalence and explained the conditions that would need to be met for them to support its development.

A relatively smaller number expressed concerns regarding potential negative consequences with some expressly objecting to interfering with nature at all. These concerns were usually strongly worded and definitive. Some people expressed caution or doubt in the technology by mentioning past mistakes in biological control (such as the introduction of cane toads in Australia), highlighting the need to carefully approach the development of this technology and to ensure that there is an additional focus on preventing the problem (of global warming) in the first place, and considering alternative interventions. Unsurprisingly, there was a small proportion who admitted that they needed more information and/or had not formed an opinion on the technology. In addition to desiring more information, some participants also expressly requested results from scientific trials.

With an understanding of these factors, it may be possible for key stakeholders to modify how they develop and implement the technology so that it aligns with these societal priorities and expectations. Some of the main take away points for this purpose include:

- Be up-front in communicating the potential risks and uncertainties, and associated risk management strategies. If risks are unknown, then communicate the precise plan of activities that will be enacted to identify the risks, well in advance of implementation.
- Use a trusted (independent, impartial, competent and experienced) governing and regulatory body to oversee the development and implementation.
- Present simple and easy-to-understand information and explanations about the current problem, the consequences and alternative, available solutions.
- Similarly, provide simple and easy-to-understand information and explanation of how genetically engineered coral would or could work.
- Expand the program of research, moving beyond a technology-centric focus to include a broader systems problem-focussed perspective, including research specialists from all domains who are relevant to conserving the reef. This also should include a technical, social and economic appraisal of alternative practical solutions, both individually and in combination with each other.
- Conduct long-term trials in controlled conditions and communicate the results of this research to the public.

## Footnotes

1 A pop-up box also provided a definition of DNA: DNA are molecules that carry genetic instructions used in development, general functioning and reproduction in all living things.

